# Cellular protein painting for structural and binding sites analysis via lysine reactivity profiling with o-phthalaldehyde

**DOI:** 10.1101/2023.09.08.556768

**Authors:** Zhenxiang Zheng, Ya Zeng, Kunjia Lai, Bin Liao, Pengfei Li, Chris Soon Heng Tan

## Abstract

The three-dimensional structure and the molecular interaction of proteins determine their roles in many cellular processes. Chemical protein painting with protein mass spectrometry can identify changes in structural conformations and molecular interactions of proteins including their binding sites. Nevertheless, most current protein painting techniques identified protein targets and binding sites of drugs *in vitro* using cell lysate or purified protein. Here, we screened 11 membrane-permeable lysine-reactive chemical probes for intracellular covalent labeling of endogenous proteins, which reveals *ortho*-phthalaldehyde (OPA) as the most reactive probe in intracellular environment. An MS workflow was developed and coupled with a new data analysis strategy termed RAPID (Reactive Amino acid Profiling by Inverse Detection) to enhance detection sensitivity. RAPID-OPA successfully identified structural change induced by allosteric drug TEPP-46 on its target protein PKM2, and was applied to profile conformation change of the proteome occurring in cells during thermal denaturation. Application of RAPID-OPA on cells treated with geldanamycin, selumetinib, and staurosporine successfully revealed their binding sites on target proteins. Thus, RAPID-OPA for cellular protein painting permits the identification of ligand-binding sites and detection of protein structural changes occurring in cells.

**Significance Statement:** Protein painting can be used to identify changes in the three-dimensional structure and molecular interaction of proteins that govern many cellular processes but are mostly applied to cell lysate or purified protein. We identified lysine reactive probes for the intracellular labeling of endogenous proteins, and developed an MS procedure with new data analysis strategy termed RAPID-OPA to characterize the intracellular conformation change of the proteome during thermal denaturation, and identified structural change mediated by allosteric regulator TEPP-46 on target protein PKM2. Furthermore, the approach could identify ligand binding sites exemplified by labeling of target proteins in cells treated with geldanamycin, selumetinib and staurosporine. Overall, RAPID-OPA for cellular protein painting enables the detection of protein structural changes happening in cells as well as the identification of ligand-binding sites.

## Introduction

Proteins perform cellular functions through their three-dimensional structures, local conformational features (e.g. catalytic sites), and the molecular interactions in which they are involved in. Thus, investigating protein structural conformations, protein-ligand interaction, and protein-protein interaction are of great importance in cell biology and drug discovery. X-ray crystallography, nuclear magnetic resonance (NMR), and various spectroscopic techniques are commonly used to elucidate the structures of purified protein *in vitro*, thus omit the influence of native cellular environment known to affect protein structure, function and drug binding (1, 2).

Proteomics methods based on mass spectrometry (MS), such as Hydrogen-Deuterium Exchange (HDX) (3), Fast Photochemical Oxidation of Proteins (FPOP) (4-6), Covalent Protein Painting (7-9) and Chemical Cross-Linking (10-13), have been applied to study dynamic change of protein structures in cells. Chemical Cross-Linking generally suffers from low reaction stoichiometry, restricting the methodology to profiling the most abundant proteins in the proteome. HDX can be used to resolve solvent accessibility and weak hydrogen bonding, but depends on reversible covalent labeling that could compromise the detection of structural changes (14). FPOP requires elaborated instrumental and experimental setup that hinder their popular adoption for global proteome study (15, 16). Covalent Protein Painting (CPP), where solvent-accessible amino acids on proteins are covalently labeled with chemicals, is a more accessible and promising chemical proteomics strategy for intracellular studies of protein structures and conformational changes (8). Chemically modified regions or specific amino acid sites constitute a “footprint” that is affected by protein structure as well as interaction with protein, metabolites and xenobiotics such as drugs. In particular, CPP with protein mass spectrometry does not require protein purification and can analyze thousands of proteins in parallel.

The amino acids that are often labeled in common chemical proteomics strategies are cysteine and lysine, of which lysine accounts for 6% of all residues in human proteins and is one of the most represented amino acids with an average of 30 lysine sites per protein (17). Lysine is present in many functional sites such as enzyme active sites (18), protein-protein interaction interfaces (19), and post-translational modification sites. Currently, the commonly used membrane-permeable probes for labeling or cross-linking primary amines are *N*-hydroxysuccinimidyl ester (NHS ester), but suffer from poor selectivity (also react with histidine, serine, and tyrosine), poor stability and low labeling efficiency under physiological conditions (20). Thus, there is a strong need for better probes and analysis strategy to analyze intracellular changes of protein structure and interaction using CPP.

Here, we screened 11 membrane-permeable chemical probes revealing *ortho*-phthalaldehyde (OPA) with the highest reactivity intracellularly exceeding NHS esters, coherent with existing vitro studies showing that OPA is capable of amine reactions with peptides and proteins under physiological conditions, and is better than NHS esters for protein labeling (21). OPA is also used to modify proteins and peptides by intramolecular OPA-amine-thiol three-component processes for cyclization (22, 23) and OPA-amine two-component reactions for bioconjugation (24, 25). Nevertheless, while OPA is the most reactive intracellular probe identified, the number of OPA-labelled peptides identified is still limited for the practical use of CPP under physiological intracellular context. To overcome this, we develop a new data analysis strategy termed RAPID (Reactive Amino acid Profiling with Inverse Detection) coupled with an optimized MS workflow to identify intracellular changes in structures and molecular interactions of proteins. We show that RAPID-OPA can reveal global conformational changes of protein occurring in cell during thermal denaturation as well as specific conformational change induced in protein PKM2 by allosteric inhibitor TEPP-46, and identified binding sites of geldanamycin, selumetinib, and staurosporine on target proteins.

## Results and Discussion

### Validation of RAPID strategy

Lysine-reactive probes often react with proteins at low stoichiometry that necessitate enrichment to identify peptides containing the probe-modified lysine, while MS fragmentation could lead to probe cleavage generating daughter ions with unpredictable mass-to-charge ratio (m/z) (26). We hypothesized that these challenges can be circumvented by identifying lysine probes, preferably membrane-permeable, that interfere with the cleavage of proteins by trypsin at C-terminal of arginine or lysine. This will lead to decreased abundance of peptides that could have been generated otherwise by trypsin digestion of proteins. In addition, probe-modified lysine may not be amenable to further labelling in multiplexing MS analysis. Thus, by comparing unmodified peptides from control samples (without probe) to probe-treated samples with an isobaric labelling MS strategy using TMT (Tandem Mass Tag) labelling reagents, low abundant probe-modified peptides could be inferred with increased sensitivity while omitting tedious enrichment that increase experimental variability (Fig. 1A). Cell intrinsic factors, such as changes in post-translation modification, structural conformation, and interactions of proteins with metabolites, drugs and other proteins, could lead to differentiated changes in lysine reactivity that can be explored with this analysis strategy termed RAPID (Reactive Amino Acid Profiling by Inverse Detection). Accordingly, we first screened 11 chemical probes, consisting of NHS esters and diformaldehydes, incubating them with intact cells followed by cell lysis, tryptic digestion and TMT-labelling for MS analysis. We observed sample treated with membrane-permeable OPA (Fig 1B b-1) had the largest decrease in abundance of peptides identified (Fig. 1B-C). We note that OPA contains two aldehyde groups that can be condensed with an amine by Mannich reaction and/or reductive amination (Fig. S1A).

**Fig. 1.**
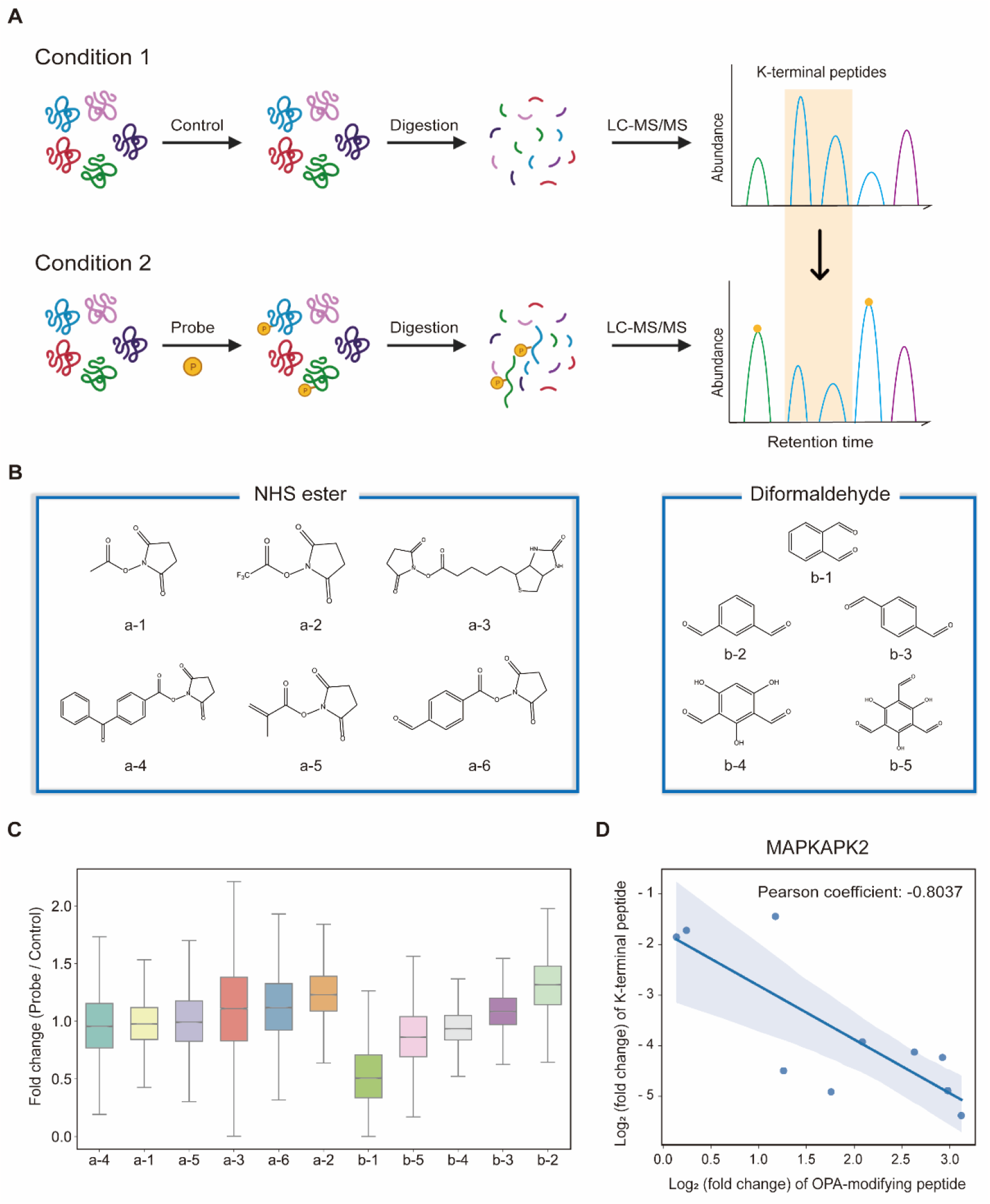
Proposal of the entire workflow for lysine reactivity profiling in complicated proteomes. (A) Principle of RAPID strategy. (B) Chemical structure of lysine reactivity probes grouped into NHS esters and diformaldehydes. (C) Boxplot visualization of abundance changes for peptides in HEK293T cell induced by each lysine reactivity probe. (D) Scatter plot shows the correlation in abundance changes of OPA-labeling peptides and their native peptides in MAPKAPK2 protein.

The abundance decrease of native peptide identified could potentially arise from direct interference of trypsin by OPA, hence steps are incorporated in our workflow to remove excess OPA prior enzymatic digestion of proteins (27). Nevertheless, to further validate that the decreased abundance of native peptides arises from increase in OPA-labelled peptides, purified MAP kinase-activated protein kinase 2 (MAPKAPK2), which contains 30 lysine residues, was incubated with OPA followed by bottom-up MS analysis where OPA is included as variable lysine modification during MS spectral search. Using purified protein at high concentration increases OPA reaction stoichiometry and the abundance of OPA-labelled peptides for MS detection. A total of 82 peptides were quantified, of which 19 peptides (23.17%) were OPA-labelled (Fig. S1B). As anticipated, the native peptides of identified OPA-labeled peptides shown a corresponding decrease in abundance (Fig. 1D and Fig. S1C). These results strongly suggested that intensity of OPA labelling can be inferred from changes in abundance of native peptides.

Next, we hypothesized that the OPA-labelling reaction follows a second-order kinetic approach (28), with the modification rate increasing in a dose and time dependent manner. To systematically optimize the labelling conditions, we first incubated HEK293T cells with different OPA concentration from 50 μM to 800 μM for 15 min reaction (Fig. 2A). Overall, we observed decreasing abundance of native peptides with increasing OPA concentration from 100 μM onward (Fig. 2B). Next, we examined how duration of incubation after different time points, namely 5, 10, and 15 min, affects labeling with 800 μM OPA. As anticipated, the abundance of native peptides decreases with time where the greatest change was observed with 15 min of OPA incubation (Fig. 2C). Overall, the concentration- and time-dependent effect of OPA suggest the decreased peptide abundance observed is largely due to reaction of lysine with OPA and demonstrated that OPA could label proteins progressively under physiological conditions.

**Fig. 2.**
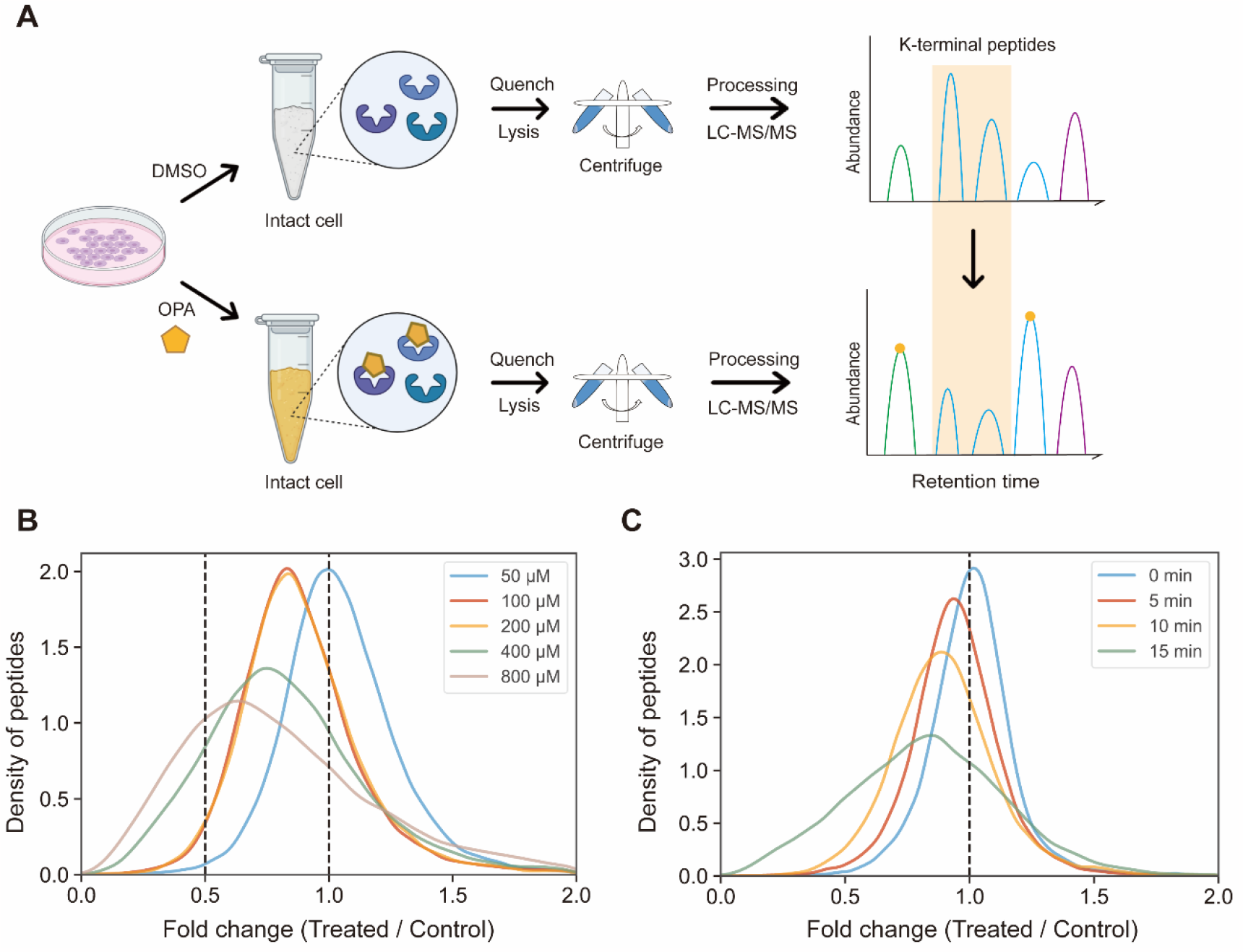
Experimental process and method optimization. (A) General workflow of RAPID-OPA. (B) Overall density plot of peptide abundance change with different concentrations of OPA. (C) Overall density plot of peptide abundance change with different labeling times of OPA.

### RAPID-OPA for monitoring proteome-wide structural changes of proteins in cells

Overall, with 15 min of 800 μM OPA incubation in HEK293T cells, 8,487 out of 63,005 native peptides (13.47%) across 6,943 proteins identified exhibited statistically significant (p-value < 0.05) decrease in abundance, while spectral search incorporating variable OPA modification could only identified 673 OPA-labeled peptides (1.07%) (Fig. 3A). In comparison, lysine reactivity was better in HEK293T cell lysates with the same OPA concentration and duration, with 31,875 (60.15%) of the 52,994 native peptides from 6,057 proteins showed a statistically significant decrease in abundance (p-value < 0.05). Spectral search identified 2,678 OPA-labeled peptides (5.05%) from cell lysate, three times more than those identified from intracellular labelling. (Fig. S2) We investigate the applicability of the RAPID-OPA workflow to study proteome-wide intracellular conformational changes of protein structure. First, using monomer structures of human proteins predicted by AlphaFold2, we observed native peptides with largest decrease in abundance after OPA labelling are over-represented in highly surface accessible lysine (Fig 3B). Nevertheless, not all peptides with highly surface accessible lysine exhibit marked decrease in abundance. As monomer structures of proteins were used in calculating solvent accessible surface area (SASA), some of these peptides might be parts of protein involved in interactions with membrane, nucleic acids, and other proteins, which could also influence protein structural conformation not predicted by AlphaFold (29). We also observed peptides with partially surface accessible lysine could exhibit marked decrease in abundance, possibly due to microenvironment of lysine with proximal amino acids that could influence its reactivity with OPA (19). Thus, changes in structural conformation of protein and lysine microenvironment could be reflected as differentiated changes to OPA labelling of lysine. To investigate this, MS data from intact cells treated with OPA at three temperature points (namely 37°C, 43°C and 49°C) were collected and analyzed (Fig. 3C). At these elevated temperatures, proteins are expected to at least partially unfold, exposing inner lysine for OPA labelling which will result in less native peptides for MS detection (Fig. 3D). Indeed, we observed further decrease in native peptide abundance when cells are incubated with OPA at denaturing temperatures (43°C and 49°C) with more than 22068 (35.03%) and 29011 (46.05%) native peptides exhibited a decrease in abundance compared to 8,487 (13.47%) at 37°C. Encouragingly, partially surface accessible lysine is enriched in peptides with the largest decrease in abundance (Fig. 3D-E and Fig. S3A-D). All in all, these results affirm that RAPID-OPA can be used to detect structural changes of proteins in cell.

**Fig. 3.**
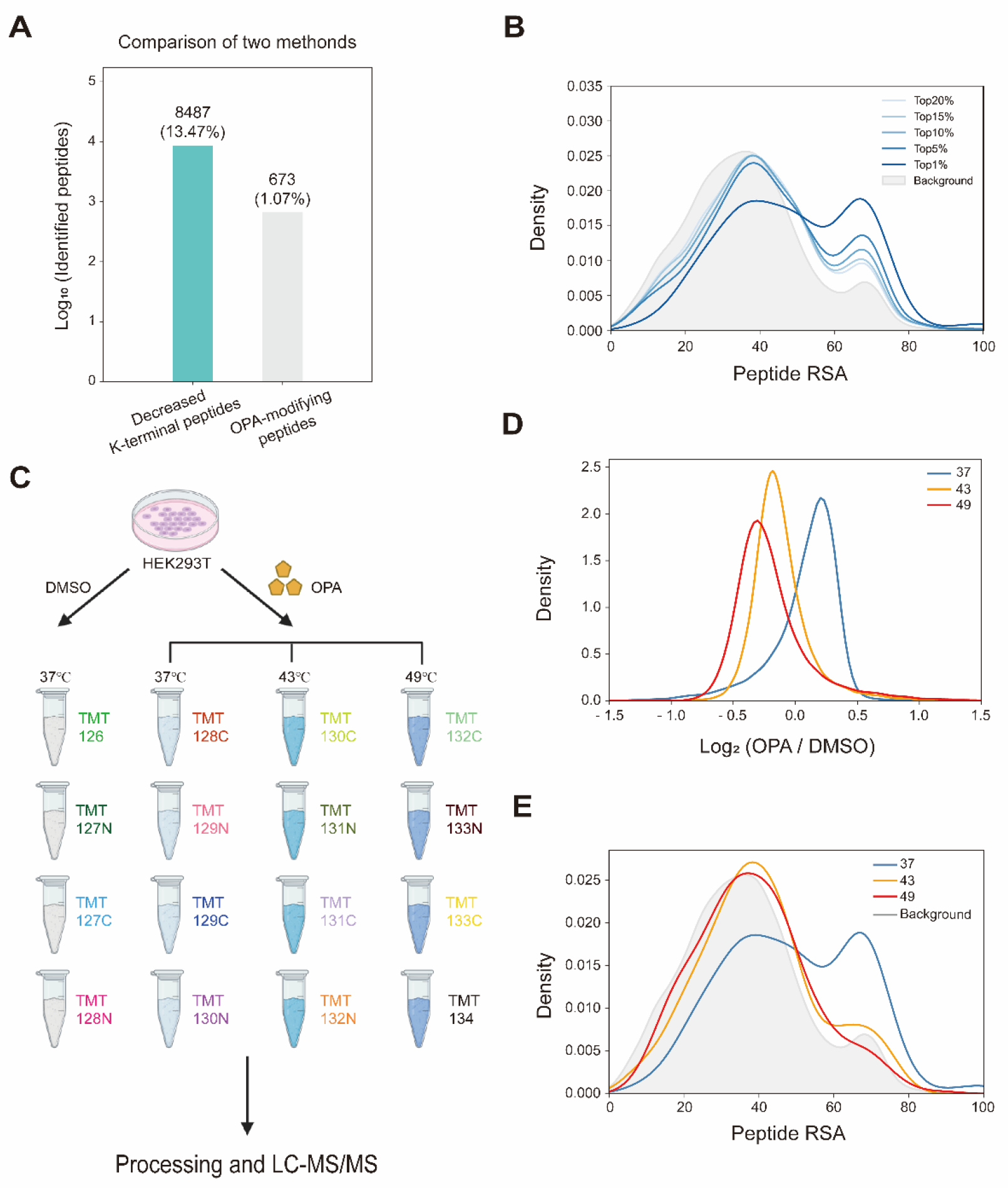
Monitoring the structural changes in proteomes. (A) Number of peptides with differentiated abundance identified in HEK293T cells. (B) RSA distribution of down-regulated peptides at 37°C. (C) Schema of samples collected. HEK293T cells are treated with DMSO/OPA and heated at 3 temperatures for RAPID-OPA profiling. (D) Distribution of changes in native peptide abundance across different temperatures. (E) RSA distribution of the top 1% down-regulated peptides.

### Identifying protein structure conformation changes with RAPID-OPA

TEPP-46 binds and activate PKM2 (pyruvate kinase M2) by promoting the tetramerization of the protein (30). A recent work treated cells with TEPP-46 followed chemical labelling after cell lysis with isotope-coded formaldehyde (CD2O) to identify structural conformation changes induced in PMK2 (31). To further validate the use of RAPID-OPA to study conformational changes in protein structures directly in cells, HEK293T cells were treated with TEPP-46 for 20 min, followed by RAPID-OPA where proteins were labelled with OPA before cell lysis and analyzed with MS in PRM mode (Fig. 4A). Four biological replicates were performed and processed for MS analysis with a single TMT set. Overall, we detected 30 native peptides of PKM2 protein, of which 3 peptides exhibit a statistical significance decrease in abundance in samples incubated with OPA. However, in cells treated with TEPP-46, the decrease in abundance due to the formation of tetramers induced by the drug is significantly muted for native peptide [K^422^].CCSGAIIVLTK^433^.[S] of PKM2 protein suggesting OPA-labelling of either the preceding Lys ^422^ or the Lys ^433^ within the peptide had been impeded (Fig.4B-C). Both Lys ^422^ and Lys ^433^ are exposed in monomer PKM2 (Fig.4E) but Lys ^422^ are lysine found on binding interface of tetramer PKM2 (Fig.4D), where the RSA of Lys^422^ and Lys^433^ in tetrameric PKM2 is less than 15% but over 90% in monomer PKM2 (Fig. 4E). As PKM2 changes from a monomer to tetramer upon treatment with TEPP-46, this result further validated the use of RAPID-OPA for studying changes in structural conformation and molecular interaction of proteins.

**Fig. 4.**
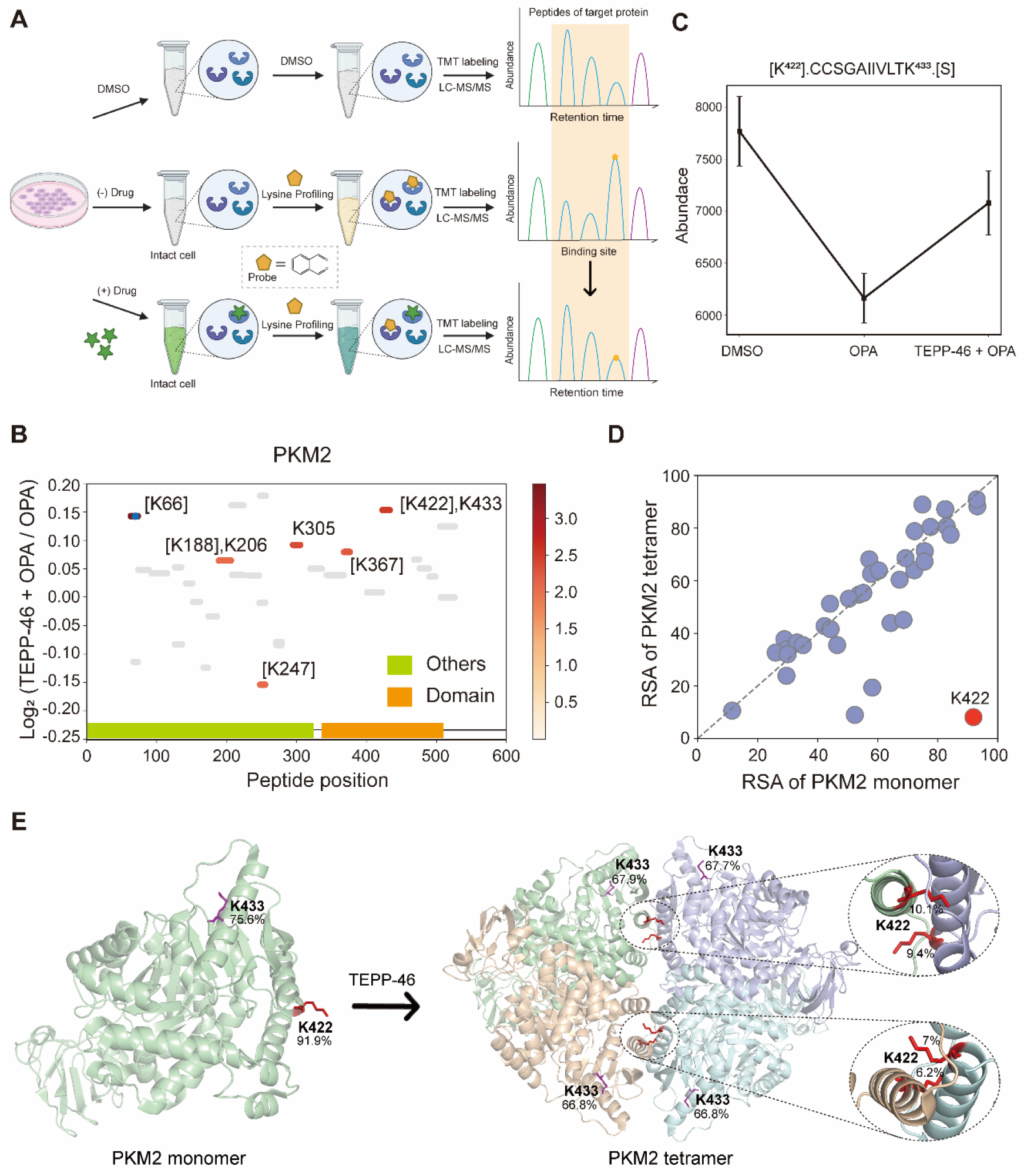
Interrogation of the allosteric proteins PKM2 induced by TEPP-46. (A) Schema of samples collected. HEK293T cells are treated with drug before OPA. (B) Diagram shows the locations and labeling reactivity shifts of identified peptides in PKM2. (C) Abundance of peptides [K^422^].CCSGAIVLTK^433^.[S] under different conditions. (D) Scatter plot shows the comparison of lysine RSA in monomer PKM2 and tetrameric PKM2. (E) RSA of K^422^and K^433^ in tetrameric PKM2 and monomer PKM2. The four PKM2 monomers and K^422^ are represented in cartoon mode. The red square is the binding region of the PKM2 protein and the drug TEPP-46.

### Interrogation of the target proteins and binding sites of geldanamycin

Next, we used geldanamycin to assess applicability of RAPID-OPA to study protein binding sites of drugs and chemicals. Geldanamycin (GA) is a natural heat shock protein 90 (Hsp90) inhibitor that specifically binds to ATP binding sites in the N-terminal domain of Hsp90 (32). RAPID-OPA was applied on HEK293T cells treated with geldanamycin with good reproducibility in peptide quantification observed among four technical replicates performed (Fig. S4A-C). The analysis identified 54 peptides and 59 peptides from known HSP90AA1 and HSP90AB1 target proteins of geldanamycin. Perhaps due to the high abundance of HSP90AA1 and HSP90AB1, their OPA-labelled peptides were identified and exhibited significant changes. Specifically, the peptides with significant lysine residues shifts in reactivity were 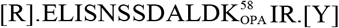 and [R].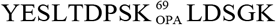 .[E] for HSP90AA1, and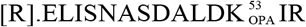 .[Y] and 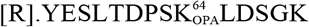 .[E] and for HSP90AB1 (Fig. 5A-B), all which were located within or near the known binding regions of geldanamycin in these target proteins (27). For HSP90AA1, Lys^58^ was located directly in the interaction region, while Lys^69^ was located near the interaction region (Fig. 5C). The abundance of these OPA-labelled peptides is significantly lower in drug-treated group suggesting OPA-labelling of either the preceding Lys^58^, Lys^69^, Lys^53^ or Lys^64^ within the binding site had been hindered by occupation of geldanamycin (Fig. 5D-E). In addition, both Lys^100^ of HSP90AA1 and Lys^95^ of HSP90AB1 were contained in native peptides, which were located in the domain with significant changes and are near the interaction region suggesting that conformational changes occurred after drug treatment (Fig. 5A-C). Thus, RAPID-OPA could also delineate regions of ligand binding on target proteins.

**Fig. 5.**
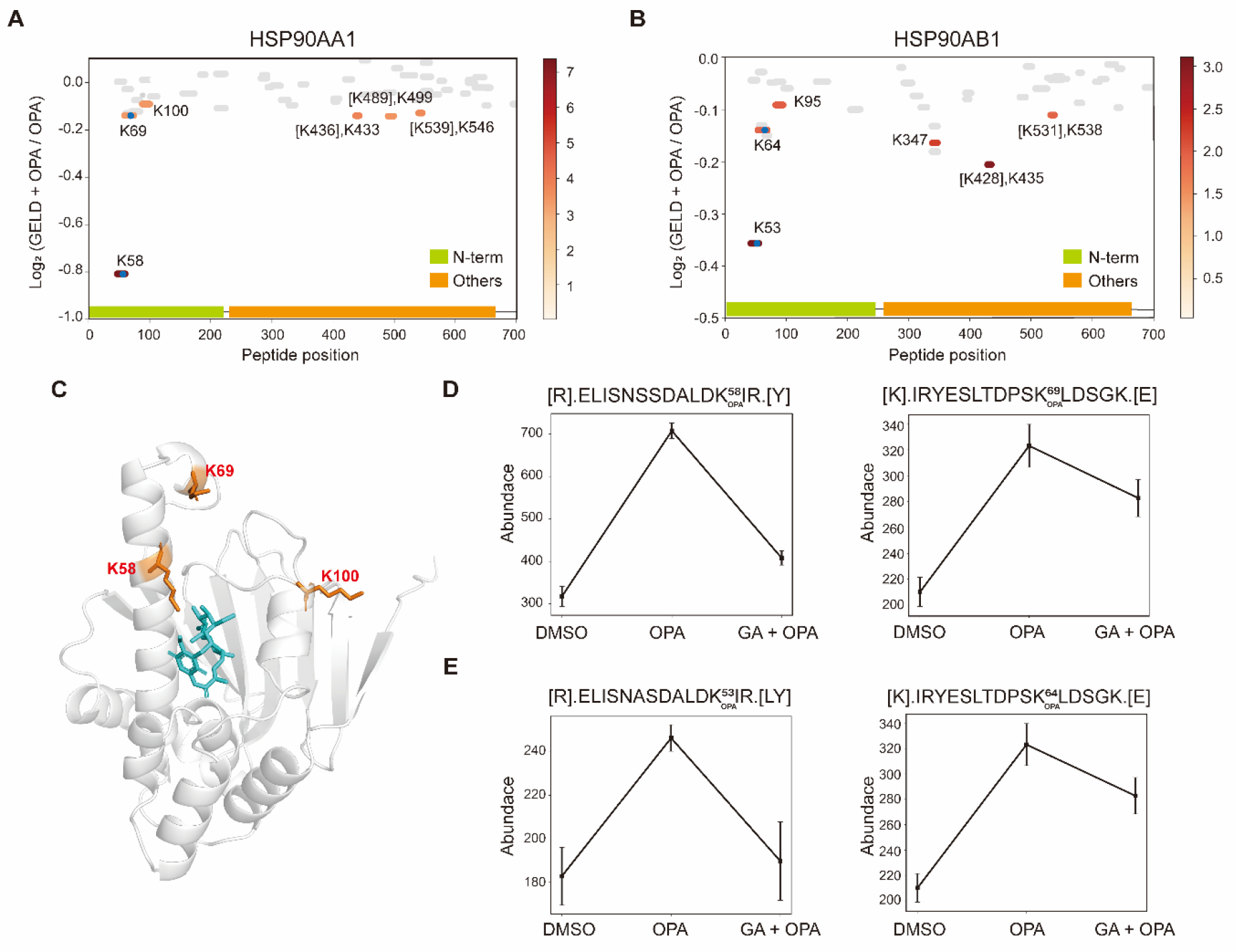
Interrogation of the target proteins and binding sites of geldanamycin. (A-B) Diagram shows the locations and labeling reactivity shifts of identified peptides in HSP90AA1 and HSP90AB1. The blue dot means OPA-label. (C) Co-crystallization result of HSP90AA1(PDB ID:1YET). (D) Abundance of peptides [R].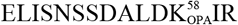 .[Y] and [K].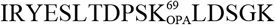 .[E] in HSP90AA1. (E) Abundance of peptides[R].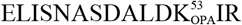 .[Y] and [R].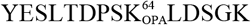 .[E] in HSP90AB1.

### Interrogation of kinase inhibitors selumetinib and staurosporine with RAPID-OPA

Next, we applied RAPID-OPA to interrogate binding sites of kinase inhibitor selumetinib on target proteins in HEK293T cells as described above in PRM mode. Selumetinib (SMT) is a selective, non-ATP-competitive, allosteric inhibitor of MEK1/2, which inhibits the RAS/RAF/MEK/ERK pathway. The result identified 11 peptides and 5 peptides from known target proteins MEK1 and MEK2, respectively. Among these, 4 and 1 significant peptides with lysine reactivity shifts of MEK1 and MEK2 respectively were located on the kinase domain (Fig. 6A-B). For MEK1, Lys^97^ was located directly in the drug-protein binding site, while other significant lysines were located near interaction region (Fig. 6C). These results show that RAPID-OPA can be applied to study the effect of kinase inhibitors on kinase domain in kinase protein.

**Fig. 6.**
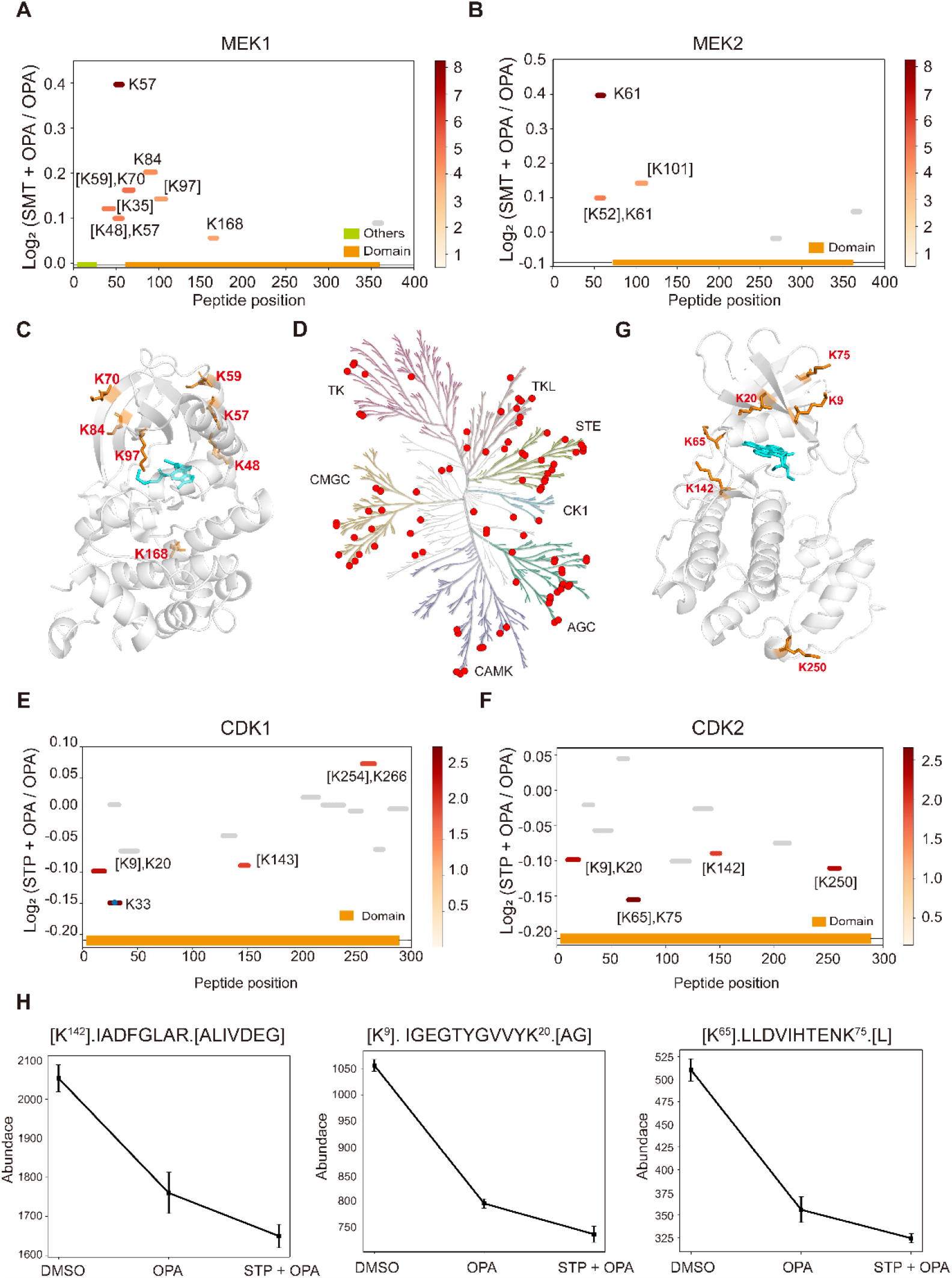
Interrogation of kinase inhibitors selumetinib and staurosporine with RAPID-OPA. (A-B) Diagram shows the locations and labeling reactivity shifts of identified peptides in MEK1 and MEK2. (C) Co-crystallization result of MEK1(PDB ID:7M0T). (D) Distribution of identified kinase proteins of staurosporine in HEK293T cells. (E-F) Diagram shows the locations and labeling reactivity shifts of identified peptides in CDK1 and CDK2. The blue dot means OPA-label. (G) Co-crystallization result of CDK2 (PDB ID:4ERW). (H) Abundance of peptides [K^142^].IADFGLAR.[ALIVDEG], [K^9^]. IGEGTYGVVYK^20^.[AG] and [K^65^].LLDVIHTENK^75^.[L] in CDK2.

To further validate the feasibility of our method, we performed a proteome-wide scan for changes in lysine reactivity in HEK293T treated with Staurosporine (STP) which is a potent, ATP-competitive, non-selective broad-spectrum inhibitor of protein kinases that binds to their ATP binding pockets. A total of 4,173 peptides with significant lysine reactivity shift were identified, with 183 (4.39%) peptides from 100 protein kinases and 467 (11.19%) peptides from 153 ATP-binding proteins (Fig. S5A-C), which are over-represented among proteins with significance changes in peptide abundance (p-values < 0.05), consistent with the known binding proteins of the staurosporine (Fig. 6D). Of the peptides from kinases that showed significant changes in abundance, they were predominantly located at the kinase domain of these kinase proteins (Fig. S5D), further demonstrating the feasibility of our approach to identify drug targets and resolve binding sites and proteome-wide level. For example, each four peptides of target kinase protein CDK1 and CDK2 with significant changes were located at the protein kinase domain and ATP binding region (Fig. 6E-F). After further analysis, we also found that Lys^9^, Lys^20^, Lys^65^, Lys^75^ and Lys^142^ with significant shift changes were located within or near the known binding regions of staurosporine in target proteins CDK2. (Fig. 6G). For CDK2, the abundance of peptides [K^142^].IADFGLAR.[ALIVDEG], [K^9^]. IGEGTYGVVYK^20^.[AG] and [K^65^].LLDVIHTENK^75^.[L] which contain above lysine was lower in drug-treated group suggesting that CDK2 protein undergoes a conformational change that exposes more lysine of the domain, which is labeled by OPA after drug treatment. In addition, staurosporine also caused significant changes in the reactivity of some lysine residues at other structural domains of kinase proteins (Fig. S5D). This indicate that the binding of staurosporine to the structural domain of these kinase may induce a conformational change. All of the above findings suggested that lysine reactivity analysis method worked well in drug-induced conformational changes of target proteins in complex proteomes and even in screening drug targets and their binding sites.

## Conclusion

In brief, we developed an intracellular chemical covalent labeling method based on lysine reactive shift coupled with a new data analysis strategy RAPID to analyze the intracellular conformational changes of proteins on a large scale. RAPID-OPA is able to detect global structural changes in proteins induced by elevated temperature. In addition, four drugs were used to benchmark the applicability of RAPID-OPA to study specific conformational changes and to identify ligand-binding sites. Thus, RAPID-OPA can be used to study intracellular protein conformational changes and for the identification of drug targets and their binding sites on proteins. We also envision RAPID to be readily deployable tools for biologist to study structural dynamics of the proteomics and molecular interactions for various cellular processes and guide mechanistic studies of bioactive compounds.

## Materials and Methods

### Reagents and Cell Culture

TEPP-46, geldanamycin, selumetinib, and staurosporine were purchased from MedChem Express. All the compounds were purchased from bidepharm (China). All the compounds were made up as solutions in DMSO (100x).HEK293T cells were cultured in DMEM medium (Gibco) supplemented with 10% FBS (PAN) and 1% penicillin-streptomycin (Gibco) at 37°, and 5% CO_2_ in a humidified environment.

### In vivo covalent small molecules for lysine reactivity

HEK293T cells were treated with 800 μM individual probes or corresponding concentration of vehicle, dimethyl sulfoxide (DMSO), incubated at 37°C for 15 min followed by addition of glycine solution to a final concentration of 2 mM with 10 min incubation to quench the reaction. Then, each mixture subsequently subjected to (2x) lysis buffer containing a final concentration of 100 mM HEPES (pH 7.5), 20 mM MgCl_2_, 10 mM β-Glycerophosphate (sodium salt hydrate), 2 mM Tris(2-carboxyethyl) phosphine hydrochloride (TCEP), 0.2 mM Sodium orthovanadate,0.2%(w/v) n-dodecyl β-D-maltoside (DDM), and EDTA-free protease inhibitor (Sigma-Aldrich, USA). Cell suspension was subjected to five times flash-freezing in liquid nitrogen and rapid thawing in water to facilitate cell lysis. After centrifugation at 21,000 g for 20 min at 4 °C, the supernatant was transferred to a new tube and the protein concentration was measured by the BCA assay kit (Thermo Fisher Scientific, USA).

### RAPID-OPA analysis in purified protein

The MAP kinase-activated protein kinase 2 (MAPKAPK2) samples were prepared at 2 mg/mL with PBS (pH 7.4). Then, the MAPKAPK2 samples were subjected to labeling and quench step as above. After centrifugation at 21,000 g for 20 min at 4 °C, the supernatant was transferred to a new tube and the protein concentration was measured by the BCA assay kit.

### Optimize RAPID-OPA labeling-time

HEK293T cells were treated either with 800 μM OPA or corresponding concentration of DMSO, incubated at 37°C for different duration (15 min, 10 min, 5 min, 0 min) followed by addition of glycine solution to a final concentration of 2 mM with 10 min incubation to quench the reaction. The following steps were same as above.

### Optimize RAPID-OPA reaction concentration

HEK293T cells were treated either with different concentration (50 μM, 100 μM, 200 μM, 400 μM, 800 μM) of OPA or corresponding concentration of DMSO, incubated at 37°C for 15 min. The following steps were same as above.

### Heat stimulation treatment

HEK293T cells were treated either with 800 μM OPA or corresponding concentration of DMSO and heated in parallel in a PCR (VWR, Doppio Gradient) block for 15 min to the three temperatures (37°C, 43°C, 49°C). The following steps were same as above.

### Profiling of lysine reactivity

HEK293T cells were treated with individual drugs (20 μM for TEPP-46, selumetinib, staurosporine and 100 μM for geldanamycin) and incubated at 37°C for 20 min. Next, the cells were incubated with either 800 μM OPA or corresponding concentration of DMSO at 37°C for 15 min. The following steps were same as above.

### In-solution digestion and TMT label

Each sample had 100 μg proteins in 60 μl 1x lysis buffer followed by reduced with 10 mM DTT in 95 °C for 10 min, and alkylated with 30 mM IAA for 30 min at room temperature. Then, protein was purified by methanol-chloroform precipitation (methanol/chloroform/water = 4:1:2). After lyophilized to dryness, protein pellet was resuspended in 8 M urea buffer and diluted to 2 M by adding 50 mM ammonium bicarbonate (ABC). Proteins were digested overnight at 37°C with trypsin (Promega) and Lys-C (Wako,125-05061) (trypsin/Lys-C/protein = 2:1:100 = w/w/w). The peptide concentration was measured by the Nanodrop (Thermo Fisher Scientific, USA). The samples of different compounds, RAPID-OPA labeling-time, and different concentration which no need to label TMT were directly desalted with C18 disk (3 M Empore, U.S.A.) followed by dried vacuum. The other tryptic digests samples were desalted with C18 disk and labeled with TMTpro reagent (Thermo Fisher Scientific, USA) followed by dried vacuum. The TMT-mixed sample is then fractionated into six fractions by a stepwise increasing gradient of ACN. Finally, the eluted peptide samples were lyophilized to dryness and redissolved in 0.1% (v/v) formic acid in water for nano-LC-MS/MS analysis.

### LFQ-RAPID analysis

Samples that were not labeled with TMT were subjected to analysis using a Q-Exactive instrument coupled with an Easy-nLC 1200 system (Thermo Fisher Scientific). The analytical column, consisting of an integrated spray tip (100 μm i.d. x 20 cm), was packed with 1.9 μm/120 Å ReproSil-Pur C18 beads (Dr. Maisch GmbH) for peptide separation at a flow rate of 250 nL/min. The samples were directly loaded onto the column with buffer A [0.1% (v/v) FA in water]. An 80 min gradient separation was performed as follows: 4%-8% buffer B [90% (v/v) ACN in buffer A] over 2 min, 8%-25% buffer B for 50 min, 25%-40% buffer B for 10 min, 40%-97% buffer B over 2 min, followed by 16 min wash with 97% buffer B. Peptides were detected on the Q-Exactive instrument, utilizing full MS scans on the Orbitrap mass analyzer with a resolution of 70,000, and top 10 MS2 scans at a resolution of 35,000. HCD fragmentation was performed with a normalized collision energy (NCE) of 32, and a dynamic exclusion time of 30 seconds was applied. Raw files were searched using Proteome Discoverer (PD) software (Version 2.4, Thermo Fisher Scientific) against the human proteome fasta database (Uniprot,20,376 entries, downloaded on May 03, 2022). The maximum missed cleavage for trypsin digestion was set to 2. The mass tolerance for peptide precursors was 10 ppm and the mass tolerance for fragment ions was 0.02 Da. OPA modification (+116.1062 Da), oxidation (+15.995 Da) and deamidation (+0.984 Da) of lysine residues were selected as variable modifications. FDR control for protein and peptide is 1% at strict level and 5% at relaxed level.

### Multiplexed LC-RAPID analysis

TMT-labeled samples were diluted with 0.1% FA prior to separation on 20 cm x 100 μm EASY-Spray C18 LC column with a 135 min gradient on an UltiMate 3000 HPLC system (Thermo Fisher Scientific). The mobile phase was solvent A (0.5% acetic acid in water) and solvent B (80% ACN, 0.5 % acetic acid in water). MS data were acquired by Orbitrap Exploris 480 mass spectrometer (Thermo Fisher Scientific): MS1 scan resolution was 60,000 with the *m/z* range of 350-1,200, AGC target set as standard and the maximum injection time is 45 ms. MS2 scan was collected at first mass(*m/z*) 110 and Orbitrap resolution was 30,000 using turbo TMT with HCD collision energies at 38. Raw data was also searched by PD, add TMT pro modification (+304.207 Da) as variable modifications and others setups were the same.

### TMT-PRM analysis

For PRM method, the LC-MS/MS instruments and chromatographic setups were identical to multiplexed LC-RAPID method. The PRM scan mode: MS scan mode was set at the maximum injection time of 50 ms, AGC target set as custom at the resolution of 60,000. MS2 scan mode was set at the maximum injection time of 100 ms, AGC target set as custom and an isolation window of 0.7 m/z at the resolution of 30,000. Then 38% normalized collision energy was set for HCD fragmentation. Targeted PKM2 and MEK precursor transitions and their corresponding m/z were extracted from the previous DDA discovery run and put into the PRM method as an inclusion list. Raw data was also searched by PD and the setups were the same as above.

### RSA prediction and data filtering

Solvent accessibility prediction was calculated using GetArea as described in Fraczkiewicz et al. (http://curie.utmb.edu/getarea.html). All human protein structures were downloaded from AlphaFold2 and imported into GetArea to get the accessibility profiles for each residue. The RSA value is the ratio of real side-chain SASA to average SASA in the tripeptide Gly-X-Gly in an ensemble of 30 random conformations. In this study, we calculated the peptide RSA value: the average RSA value of all amino acids in the peptide, which considered the solvent accessibility of entire peptide, and focused on local solvent accessibility of the lysine site(s). For any predicted protein structure by AlphaFold2, all residues were given the per-residue confidence scores (pLDDT) values, with values greater than 70 indicating confident results.

For analysis of OPA-modifying and K-terminal peptides detected in the proteomics dataset, any contaminant peptides and null signal peptides were removed, and only unique peptides are retained after deduplication by sequence. Then, all lysine residues were identified and matched to GetArea-predicted RSA and AlphaFold2 pLDDT values by UniProt accession ID and sequence, and the low-confidence protein structures (pLDDT value < 70) were not included for structure analysis.

## Data avalibilty

All the raw MS data have been uploaded onto the ProteomeXchange Consortium via the iProX partner repository with the dataset identifier PXD046150(33, 34).

## Acknowledgments

This work is supported by grants from and National Key Research and Development Program of China (No. 2021YFA1302603), Shenzhen Innovation of Science and Technology Commission (No. JCY20200109140814408) and the National Natural Science Foundation of China (Nos. 22074060, 22150610470) awarded to Chris Soon Heng Tan.

