## Supplemental Figure 1-5 for "Cellular protein painting for structural and binding sites analysis via lysine reactivity profiling with o-phthalaldehyde": Supplementary information.docx

**This PDF file includes:**

Supplementary figures 1-5

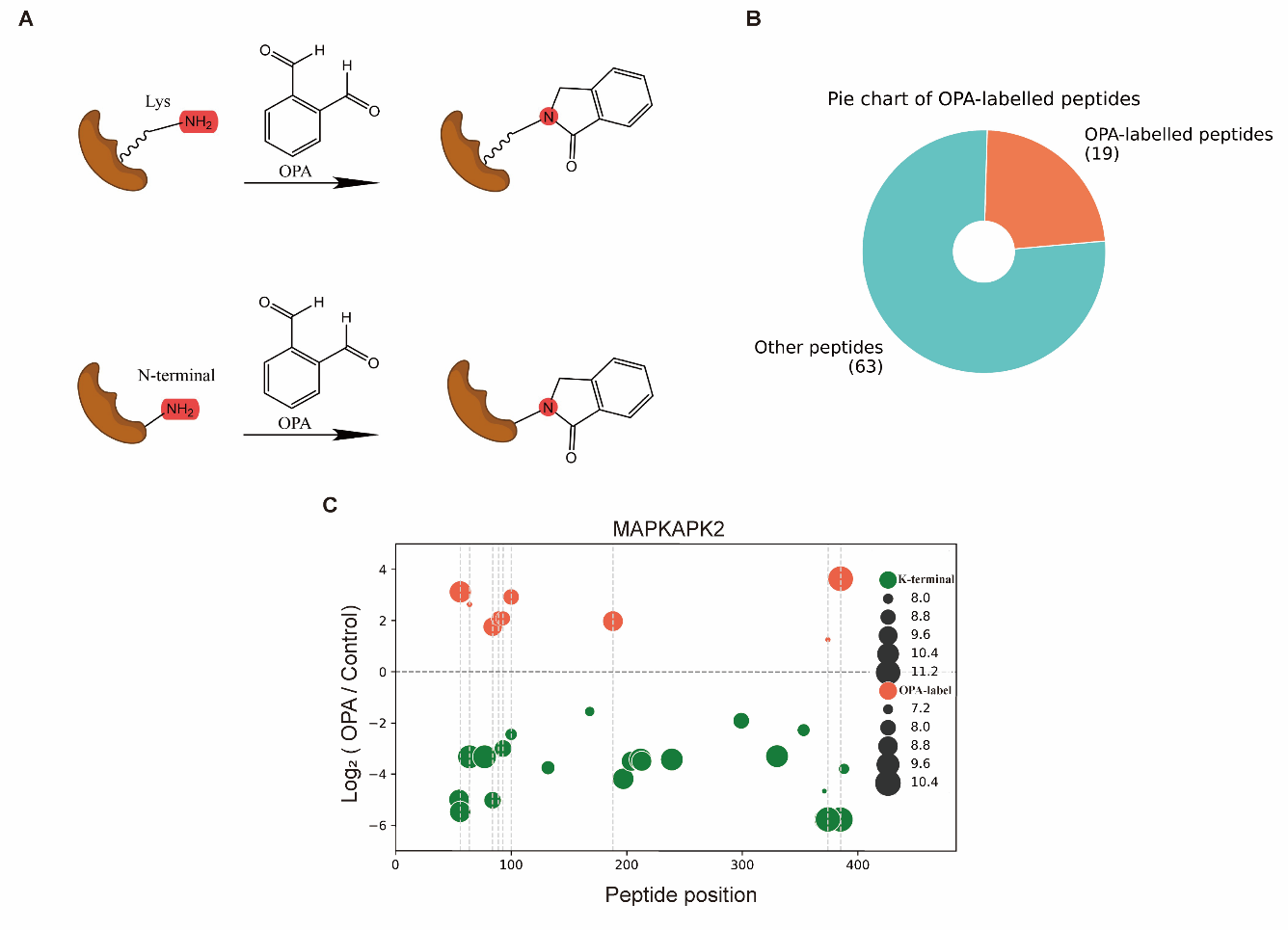
**Supplementary Figure 1-5**

**Supplementary Figure 1.** **Feasibility of RAPID method.** (A) Common strategies for peptide labeling via OPA. (B) Fraction of total quantified peptides that were liganded by OPA in MAPKAPK2 protein. (C) Bubble diagram shows the distributions and labeling reactivity shifts of identified lysine residues in MAPKAPK2 protein.

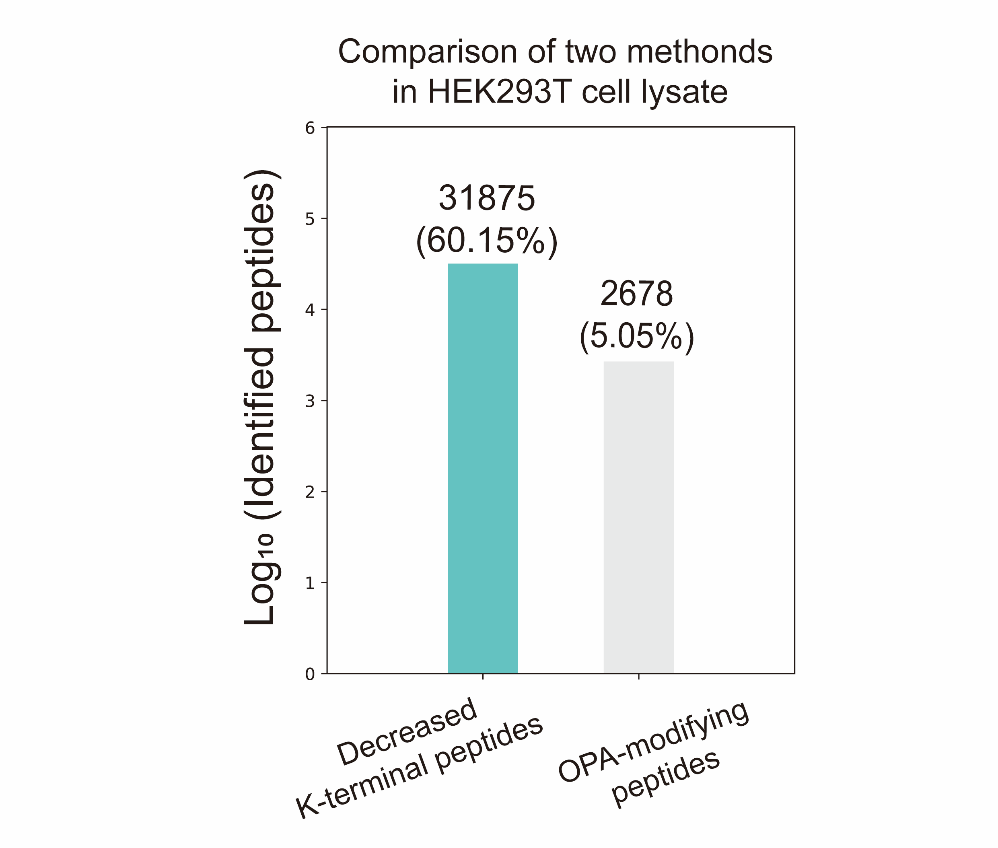

**Supplementary Figure 2.** **RAPID-OPA analysis in HEK293T cell lysate.** The number of peptides with differentiated abundance identified in HEK293T cells lysate.

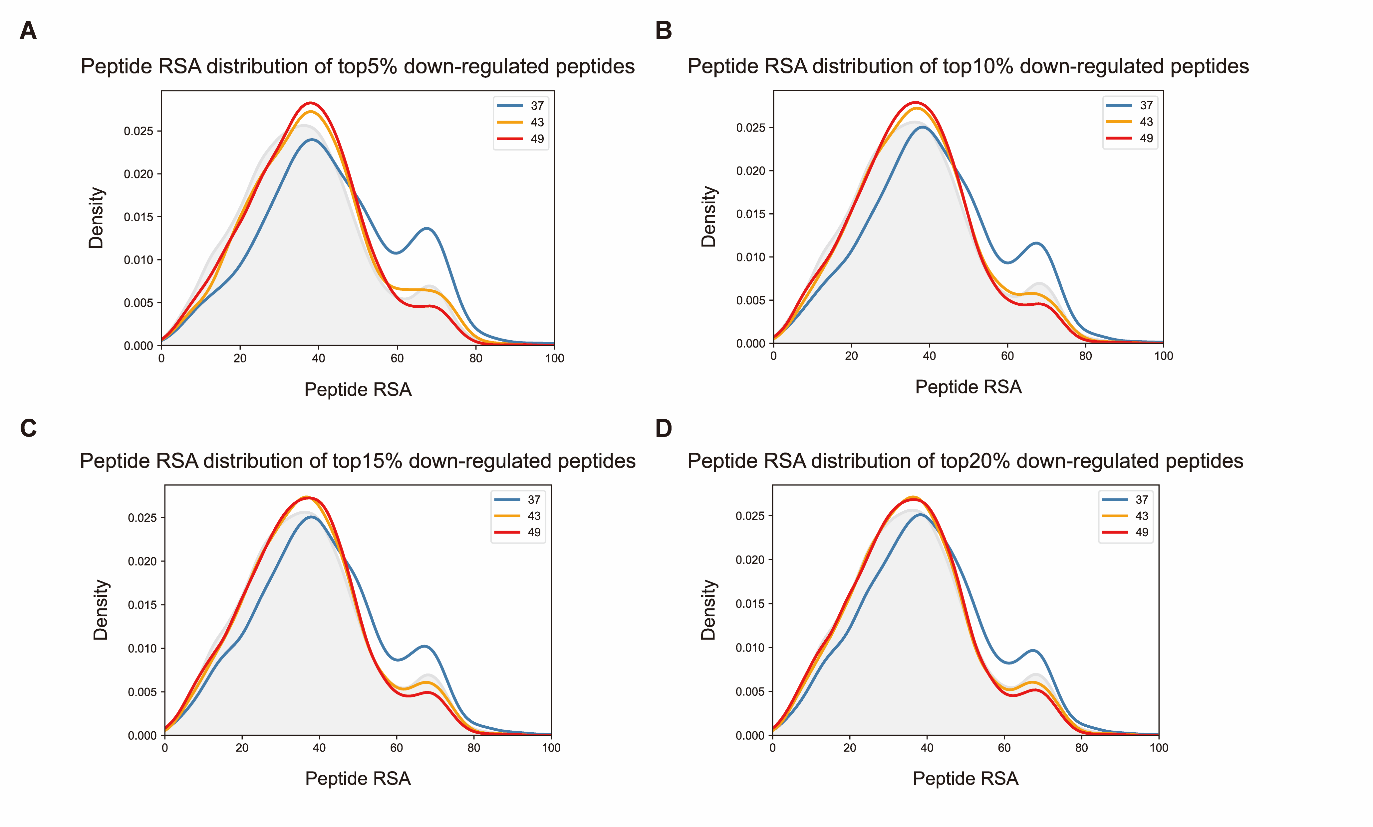

**Supplementary Figure 3.** **RSA distribution of top 5%/10%/15%/20% down-regulated peptides at 37°C, 43°C, and 49°C.** (A-D) Peptide RSA.

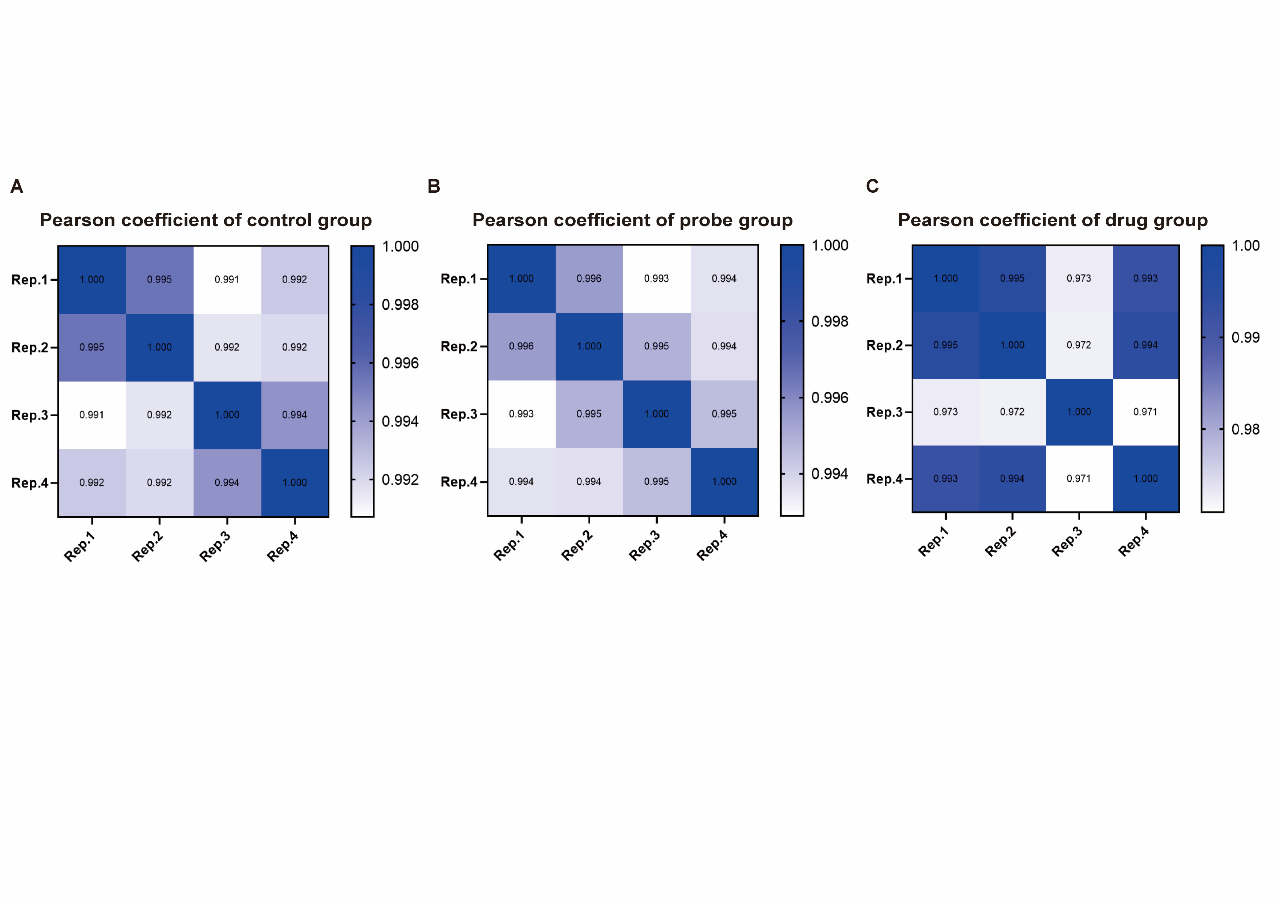

**Supplementary Figure 4.** **Pearson coefficient among four technical replicates.** (A) Pearson coefficient for four replications of the control group (treated with DMSO). (B) Pearson coefficient for four replications of the probe group (treated with OPA). (C) Pearson coefficient for four replications of the drug group (treated prior with drug before OPA incubation).

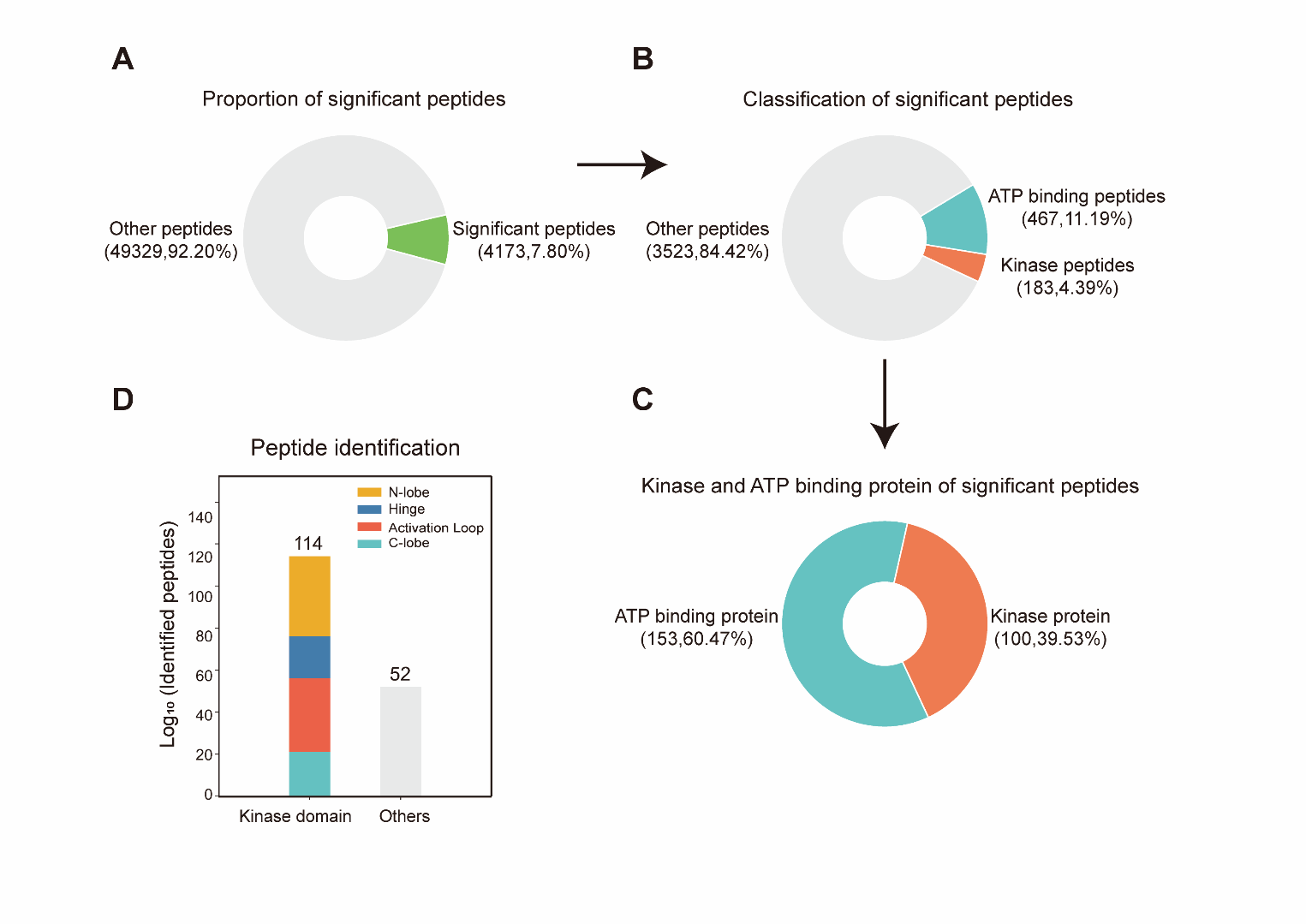

**Supplementary Figure 5.** **The effect of identifying kinase for staurosporine by RAPID-OPA.** (A) Fraction of total quantified significant peptides. (B) Fraction of total quantified kinase peptides, ATP binding peptides, and other peptides. (C) Fraction of significant peptides corresponding to kinase and ATP binding protein. (D) Bar plot visualization of the locations of the identified peptides with significantly changed labeling reactivity of kinase (adj p value < 0.05).
