## Supplementary figures and images for "Cellular protein painting for structural and binding sites analysis via lysine reactivity profiling with o-phthalaldehyde"

### supplement Figure1.pdf

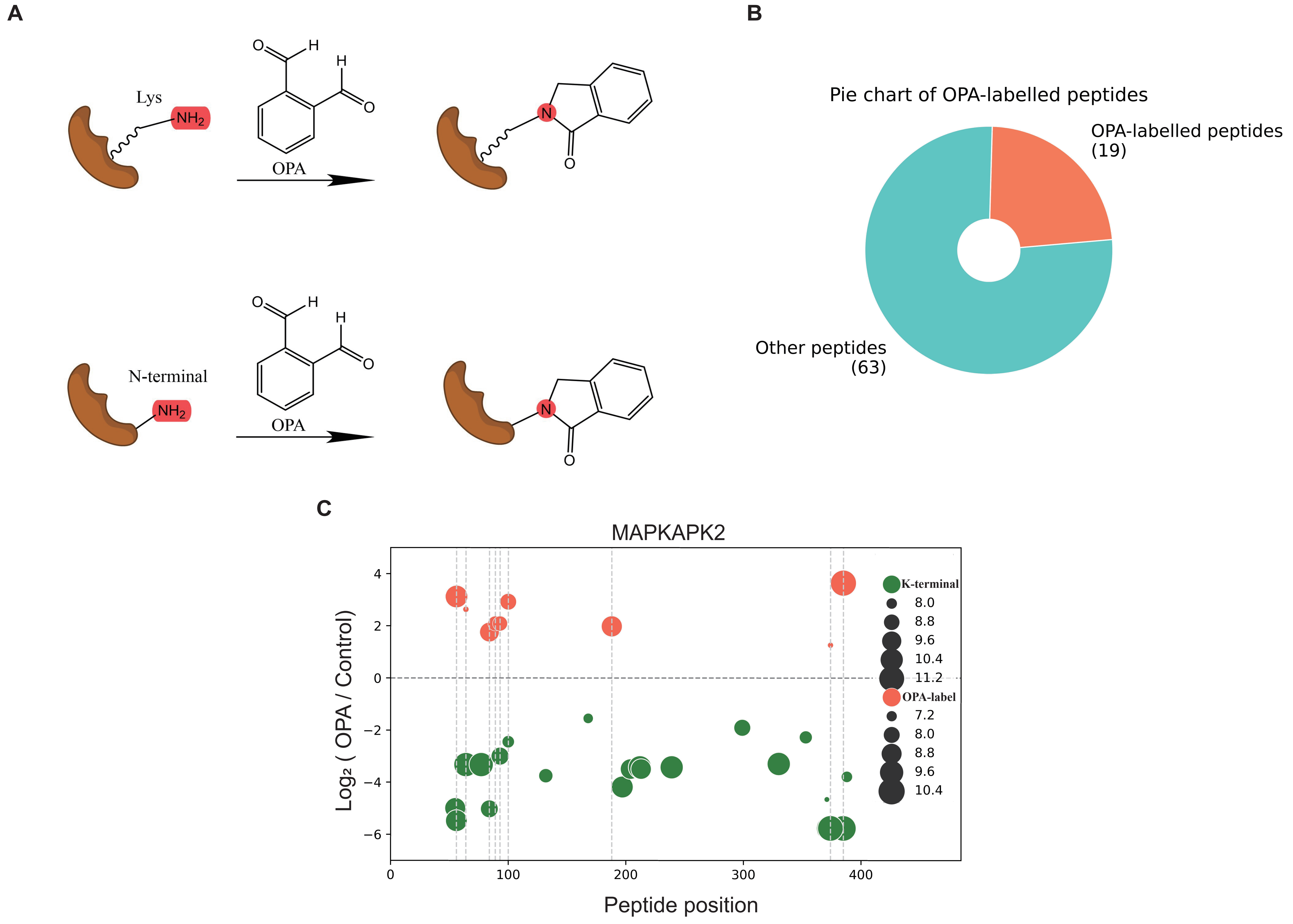

### supplement Figure2.pdf

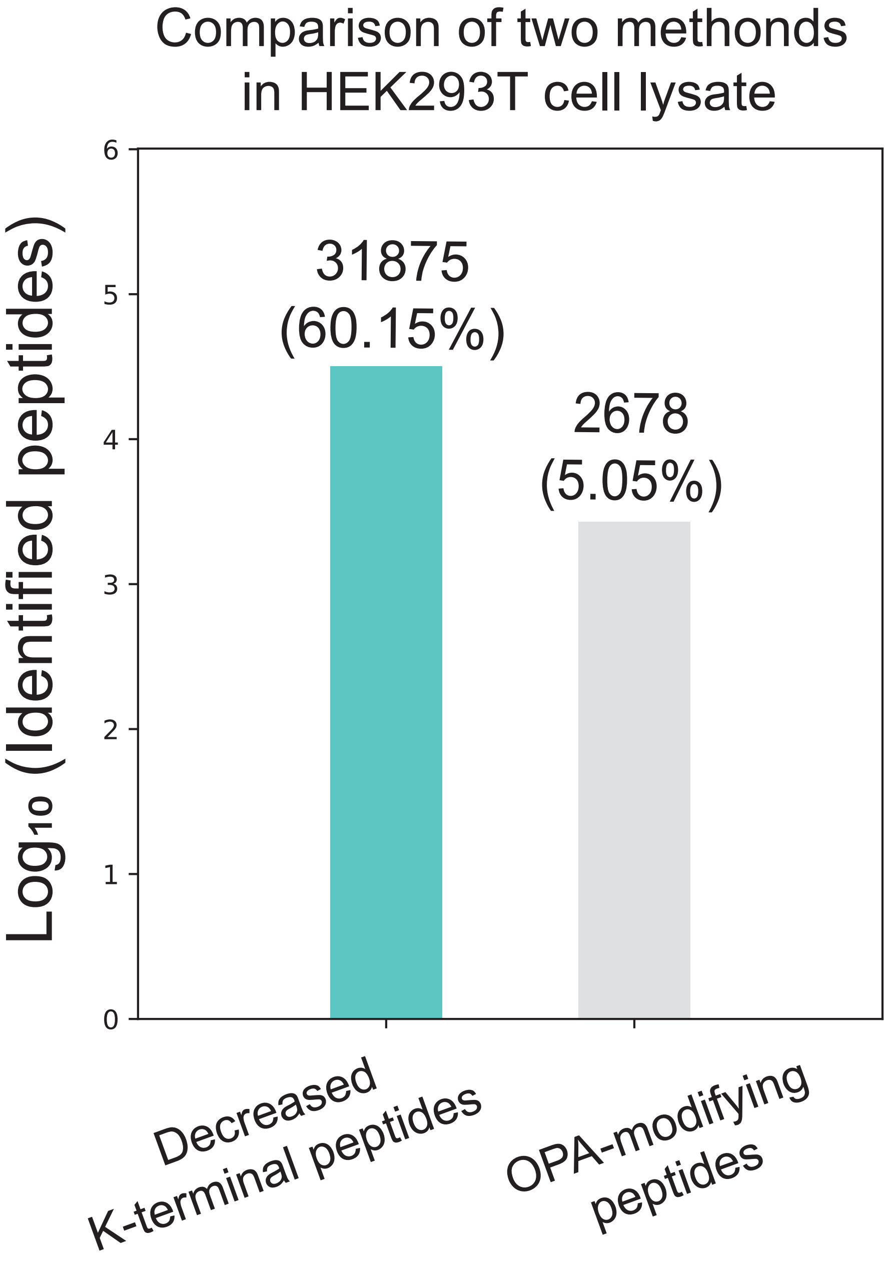

### supplement Figure3.pdf

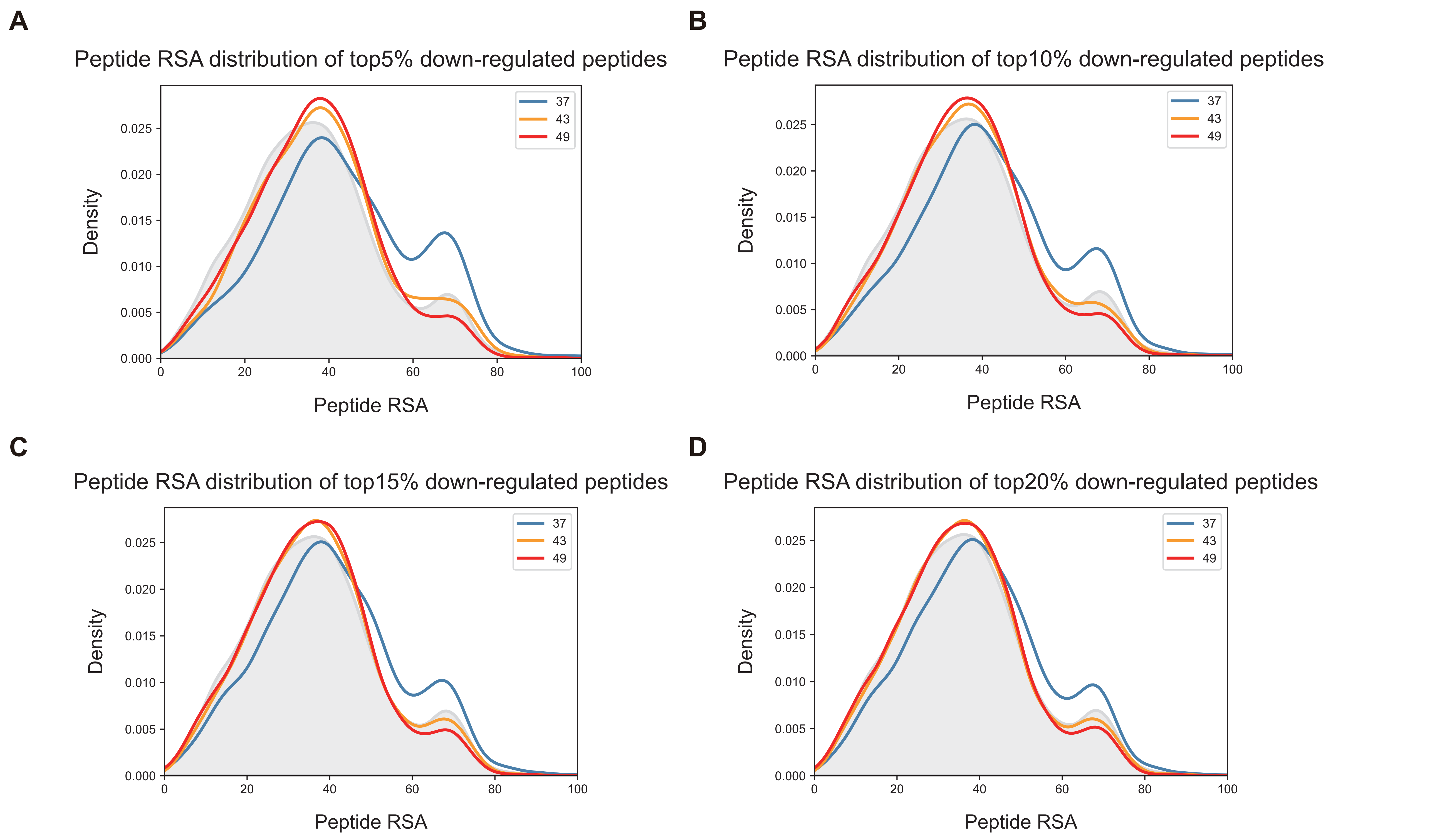

### supplement Figure4.pdf

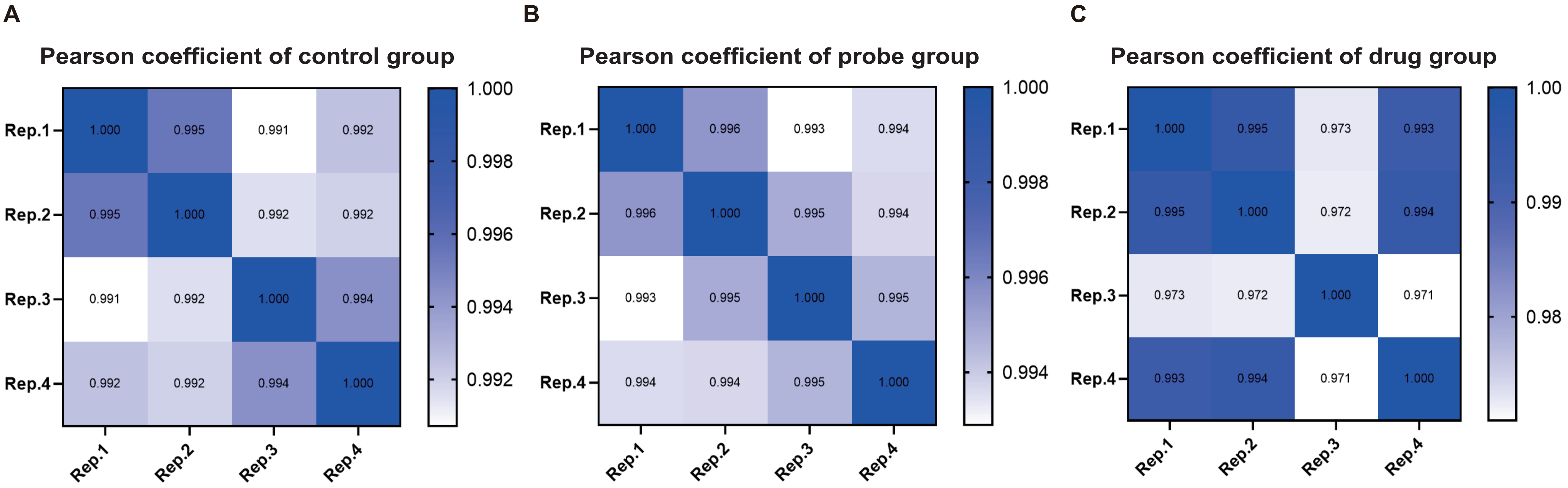

### supplement Figure5.pdf

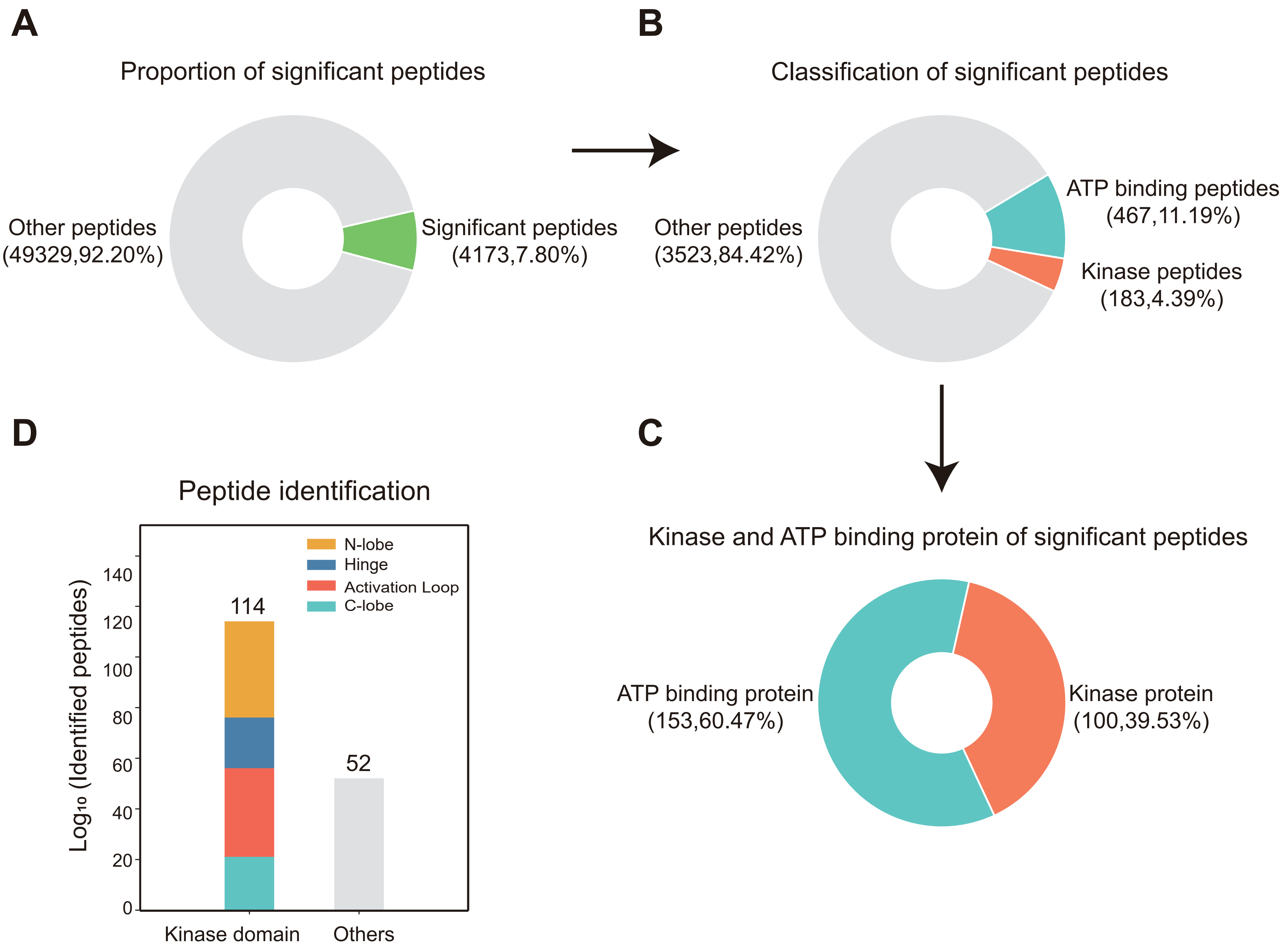
